# Genetic ablation of the TET family in retinal progenitor cells impairs photoreceptor development and leads to blindness

**DOI:** 10.1101/2024.06.19.599771

**Authors:** Galina Dvoriantchikova, Chloe Moulin, Michelle Fleishaker, Vania Almeida, Daniel Pelaez, Byron L. Lam, Dmitry Ivanov

## Abstract

Rod and cone photoreceptors are critical for vision, and their loss leads to blindness. Photoreceptors are epigenetically unique, because the promoters of the genes required for the development and function of these neurons are hypermethylated in retinal progenitor cells (RPCs) and hypomethylated in photoreceptors. However, the mechanism responsible for DNA demethylation during the differentiation of RPCs into photoreceptors and its role in photoreceptor development and function were unknown. We hypothesized that the Ten-Eleven Translocation (TET) family of dioxygenases plays a key role in this mechanism. To this end, we knocked out all TET genes in RPCs and characterized the TET-deficient and control retinas using various approaches including electron microscopy, electroretinogram (ERG) tests, RNA-seq, whole genome bisulfite sequencing (WGBS), and 5hmC-Seal. We found that genetic ablation of the TET family prevents demethylation of the promoters of genes essential for rod specification and for rod and cone maturation during the differentiation of RPCs into photoreceptors. Preservation of methylated cytosines in the promoters of these genes significantly reduced their expression, which was confirmed by western blot analysis. This impaired expression leads to the underdevelopment or complete absence of outer segments and synaptic termini in the photoreceptors of TET-deficient retinas, which results in loss of rod and cone function, as assayed by ERG. These function-deprived, underdeveloped photoreceptors die over time, which leads to blindness.

## Introduction

Rod and cone photoreceptors convert light into electrical signals, which, upon traveling to the brain, create the sensation of vision(1, 2). The loss of these neurons results in blindness. There are many signaling cascades that regulate the differentiation of photoreceptors from retinal progenitor cells (RPCs) and their subsequent maturation into functional neurons(3, 4). Impaired activity of these signaling cascades as a result of inherited mutations in the corresponding genes leads to various retinal dystrophies, such as retinitis pigmentosa, cone and cone-rod dystrophy, congenital stationary night blindness, and Leber congenital amaurosis(5-7). However, are genetic mechanisms the only ones that can influence the activity of these signaling cascades?

DNA methylation is an epigenetic mechanism that is responsible for modifying cytosines into 5-methylcytosines (5mC)(8, 9). DNA methylation in regulatory elements (e.g., promoters, enhancers) leads to reduced expression of the corresponding genes, while DNA demethylation should occur to promote gene expression(8-11). Rod and cone photoreceptors are unique cell types from an epigenetic perspective, as the promoters of many genes critical for the development and function of these neurons are hypermethylated in human and murine RPCs (12, 13). Some of these promoters were still hypermethylated in DNA isolated from murine photoreceptor precursors (14). Meanwhile, these promoters were hypomethylated and the expression of the corresponding genes was high in mature photoreceptors (12-14). However, the mechanism responsible for changes in DNA methylation patterns during RPC differentiation into photoreceptors and its significance were not understood until this study.

The patterns of methylated cytosines in DNA are established by the DNA methyltransferase (DNMT) family of enzymes that catalyze the transfer of a methyl group to the 5^th^ position in the cytosine nucleotide(8, 9). Meanwhile, the Ten-Eleven Translocation (TET) family of dioxygenases is responsible for DNA demethylation, which occurs in several steps involving the participation of additional signaling cascades and ending with the appearance of unmodified cytosines (10, 11). In this study, we examined how inactivation of the TET-dependent DNA demethylation pathway in RPCs affects the development and function of photoreceptors. We found that genetic ablation of the TET family in RPCs prevents demethylation of the promoters of key genes essential for photoreceptor development and function. This, in turn, significantly reduces their expression, preventing the development of photoreceptor outer segments and synapses. Such underdeveloped photoreceptors are deprived of function and die over time. These findings suggest the contribution of retina-specific epigenetic mechanisms to the pathogenesis of retinal dystrophies.

## Results

### Genetic ablation of the TET family in RPCs prevents proper photoreceptor development

The TET family, comprising of the *Tet1, Tet2*, and *Tet3* genes, is highly expressed at all stages of retinal development, including the earliest stage when the retina consists mostly of RPCs (**Fig. 1A, 1B**)(15). To study the role of the TET-dependent DNA demethylation pathway in photoreceptor development and function, we used transgenic animals in which exon 3 of *Tet3* and the exons encoding Tet1 and Tet2 catalytic domains were flanked by loxP sites (**Fig. 1C**)(16, 17). These animals were crossed to produce triple Tet1/Tet2/Tet3-floxed animals (TET, **Fig. 1D**). To inactivate the TET family in RPCs, we used Chx10-cre mice (Chx10/*Vsx2* is a marker of RPCs) that have been used extensively by many investigators to study retinal development (**Fig. 1D**)(18). It was shown that cre recombinase-expressing Chx10 positive RPCs differentiate into all types of retinal cells including rods and cones(18). We found that there were no significant morphological and functional differences between TET and Chx10-cre animals (**Fig. 1E-1I**).

**Fig. 1.**
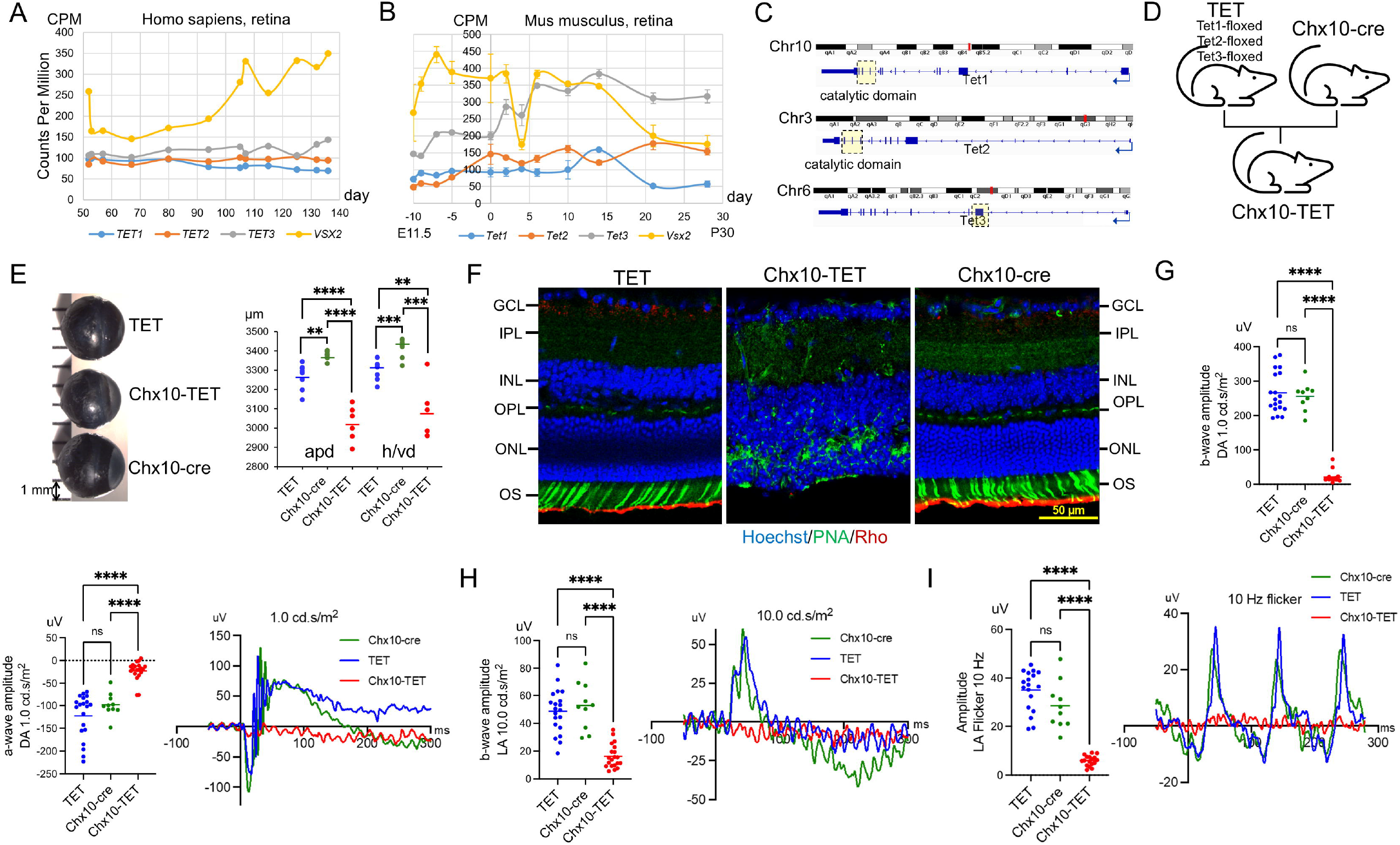
The TET-dependent DNA demethylation pathway is required for photoreceptor development and function. **(A, B)** The RNA-seq data (GSE101986 and GSE104827) indicate that members of the TET family and *Vsx2* (Chx10) gene are expressed at all stages of human (**A**) and mouse (**B**) retinal development, including the earliest stages when the retina is predominantly composed of RPCs (embryonic day (E) 11.5 in mice and human gestation days 50-60). (**C, D)** To inactivate the DNA demethylation pathway, we generated floxed TET mice in which *Tet1, Tet2*, and *Tet3* exons marked with yellow squares (**C**) were flanked by loxP sites. To inactivate this pathway directly in RPCs (**D**), we crossed TET with Chx10-cre to get Chx10-TET mice. **E)** The sizes of the eyeballs of 3-month-old (3m) Chx10-TET mice were smaller than those of TET and Chx10-cre mice of the same age (n=6-8 eyes per group; apd – anterior-posterior diameter, h/vd – horizontal/vertical diameter). **F)** While the retinas of TET and Chx10-cre mice were normally developed, the photoreceptor outer segments (OS) and the outer plexiform layer (OPL) were virtually undetectable in Chx10-TET mice (3m, n=5). **(G-I)** ERG potentials recorded from dark-adapted (rod) and light-adapted (cone) 3-month-old animals indicate that Chx10-TET mice were functionally blind. Dark-adapted (**G**) animals were subjected to a white-light flash stimulus at an intensity of 1.0 cd.s/m^2^ and frequency of 1.00 Hz. B-wave amplitudes were analyzed as the amplitude from the trough of the a-wave to the peak to the subsequent b-wave peak. The negative trough of the a-wave was analyzed at 1.0 cd.s/m^2^ intensity. Light-adapted (**H**) animals were presented with a 10 cd.s/m^2^ flash intensity ERG. Animal eyes were exposed to white light flash cycles at an intensity of 3 cd.s/m^2^ with frequencies of 10 Hz, against a background of 30 cd.s/m^2^ white light (**I**). Flicker amplitudes were measured from the N1 trough to the subsequent P1 peak. (n=10-20 eyes or 5-10 mice per group; P value <0.0001 [^****^], P value <0.001 [^***^], P value <0.01 [^**^])

However, when we crossed TET and Chx10-cre to produce Chx10-TET mice in which the RPCs lacked the activity of TET enzymes, we found significant changes in the structure and function of the photoreceptors. While the size of TET eyeballs was relatively smaller than those of Chx10-cre, the size of the eyeballs of the Chx10-TET mice was significantly smaller than those of TET and Chx10-cre (**Fig. 1E**). The retinas of Chx10-TET mice had virtually no photoreceptor outer segments (OS) and outer plexiform layers (OPL) (**Fig. 1F**). The absence of OPL resulted in the cells of the outer nuclear layer (ONL) and the inner nuclear layer (INL) mixing (**Fig. 1F**). Electroretinogram (ERG) tests conducted on dark-adapted animals showed ten times less Chx10-TET rod photoreceptor activity compared to TET and Chx10-cre (b-wave [mean±sem uV]: TET 266±14 vs. Chx10-cre 256±14 vs. Chx10-TET 21±4; a-wave: TET -123±11 vs. Chx10-cre -104±7 vs. Chx10-TET -17±3; n=10-20 eyes tested); the Chx10-TET ERG traces were almost flat (**Fig. 1G**). Similar results were obtained in the light-adapted (cone) animals (b-wave [mean±sem uV]: TET 50±3 vs. Chx10-cre 53±6 vs. Chx10-TET 14±2; flicker: TET 35±2 vs. Chx10-cre 28±3 vs. Chx10-TET 6±1; **Fig. 1H, 1I**). All these data indicate that, unlike TET and Chx10-cre animals, Chx10-TET mice were functionally blind.

### TET-deficient photoreceptors lack OS and synaptic termini, which prevents them from functioning normally, leading to retinal dystrophy

We undertook more detailed studies of Chx10-TET and TET animals since Chx10-cre animals were previously well studied and did not differ much from TET mice. To this end, we collected retinas from animals of different ages to investigate the thickness of their layers. The thickness was measured every 200 µm starting from the optic nerve head (***SI Appendix*, Figs. S1, S2**). This thickness was then averaged (**Fig. 2A, 2B**). We found that the OS and OPL (photoreceptor synapses are in this layer) were poorly developed or absent in the retinas of very young Chx10-TET animals (postnatal-day [P] 14), while OS and OPL of TET animals were well-developed (**Fig. 2A, 2B**). A more detailed analysis of our IHC data indicates the presence of underdeveloped OS of cones in P14 and 1-month-old Chx10-TET mice, but their absence in 3- and 6-month-old Chx10-TET animals (the conclusion was made based on staining with PNA, which is a marker of cone OS; **Fig. 2A, 2B**). The OS of the rods are absent in Chx10-TET mice of all ages (**Fig. 2A, 2B**). We found some differences in the thickness of the ONL (all photoreceptors are located in this layer) in P14 and 1-month-old Chx10-TET compared to TET mice. However, these differences began to become more significant with age, reaching as few as three lines of nuclei in some parts of the ONL of 6-month-old Chx10-TET animals (**Fig. 2A, 2B, *SI Appendix*, Fig. S2**). Thus, photoreceptor developmental disorders lead to their eventual death. However, we found only a few dead cells in 3-month-old Chx10-TET retinas using TUNEL assay (***SI Appendix*, Fig. S3**). These findings suggest that cell death occurs slowly in Chx10-TET retinas and surrounding cells manage to successfully phagocytose the dead cells. Therefore, we can only detect a small number of dead cells using TUNEL assay. These results are consistent with the slow retinal degeneration we observed in Chx10-TET mice (**Fig. 2A, 2B**). Meanwhile, we did not find significant changes in the thickness of the INL and inner plexiform layer (IPL) in the retinas of Chx10-TET and TET mice of all ages (**Fig. 2A, 2B, *SI Appendix*, Fig. S2**). We found that the size of Chx10-TET retinas is smaller than TET (**Fig. 2B, *SI Appendix*, Fig. S1**). These observations are consistent with the fact that Chx10-TET mice have smaller eyeballs than TET mice (**Fig. 1E**).

**Fig. 2.**
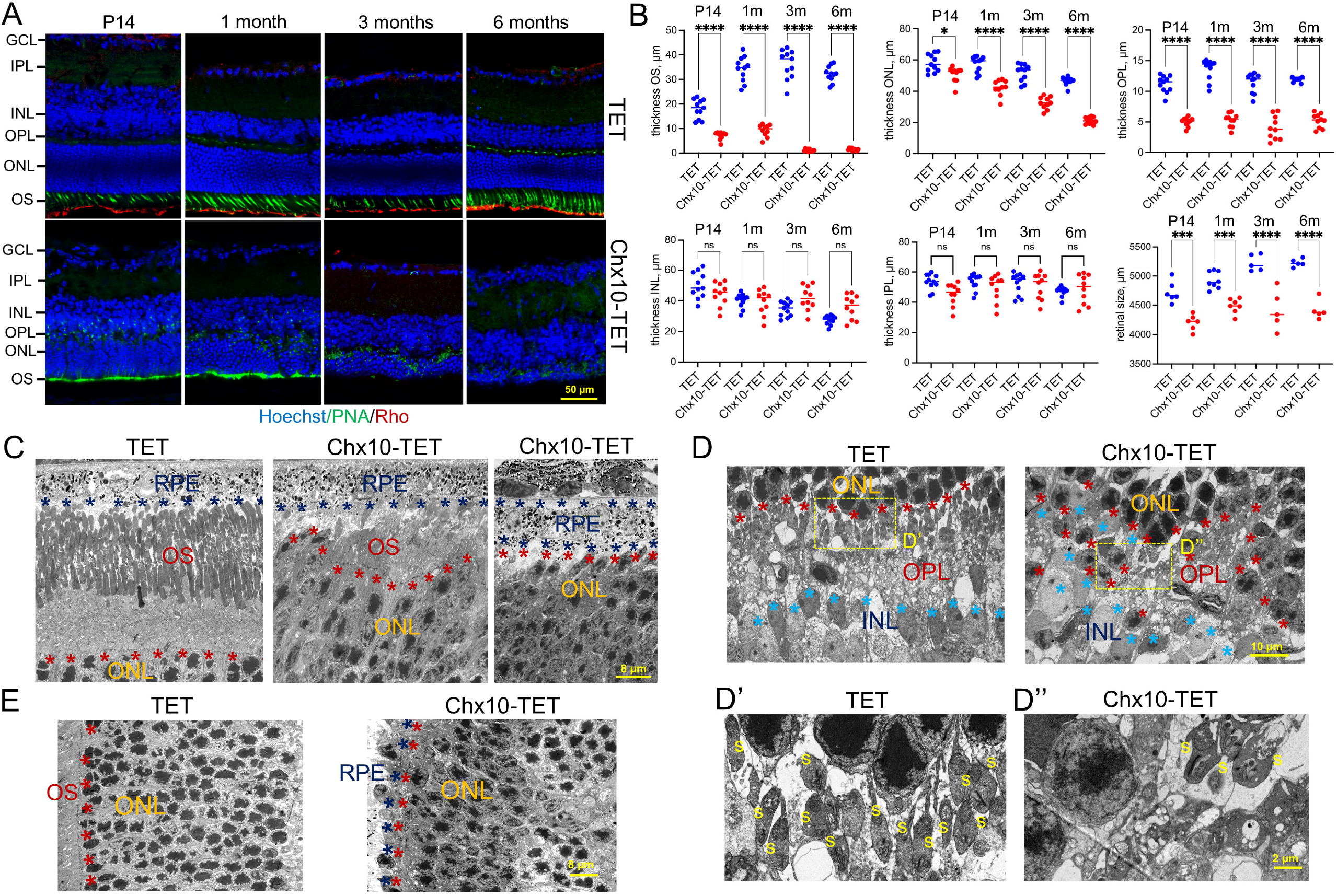
The impaired activity of the TET-dependent DNA demethylation pathway in RPCs leads to the formation of photoreceptors without OS and synaptic termini, which ends in their slow death. **A)** The representative confocal images show retinal morphology and the change in this morphology over time in TET and Chx10-TET mice (PNA is a marker of cone OS, Rho is a marker of rod OS). **B)** The results of an analysis of the thickness of the retinal layers indicate a slow degeneration of photoreceptors leading to retinal dystrophy in Chx10-TET animals. TET-deficient retinas are also smaller in size compared to TET retinas (n=5 mice per group, per age; P value <0.0001 [^****^], P value <0.001 [^***^]). **(C-E)** TEM examination of 1-month-old animals indicates underdevelopment or absence of OS **(C)** and synapses **(s, D-D’’)** in TET-deficient photoreceptors. The small number of synapses in the outer plexiform layer (OPL) leads to the fact that cells of the outer nuclear layer (ONL) and the inner nuclear layer (INL) can mix freely in Chx10-TET retinas **(D)**. Chromatin is less condensed in the nuclei of TET-deficient photoreceptors located in the ONL **(E)**. All images were collected at the same distance from the optic nerve head (800 µm). The dark blue stars limit the layer that contains retinal pigment epithelial (RPE) cells. Red stars indicate the boundary of the ONL. Blue stars indicate the boundary of the INL. n=5 retinas or 5 mice per group

To study the morphology of TET-deficient photoreceptors, we used transmission electron microscopy (TEM). TEM examination revealed well-developed OS in photoreceptors of TET animals (**Fig. 2C, *SI Appendix*, Fig. S4**). Meanwhile, the OS were undeveloped or completely absent in photoreceptors of 1-month-old Chx10-TET mice (**Fig. 2C, *SI Appendix*, Fig. S4**). We found a clear separation in TET retinas of the ONL from the INL by the OPL in which photoreceptor synapses were present. The cells of the ONL and INL were mixed in the retinas of the Chx10-TET mice; synapses are difficult to find in these retinas (**Fig. 2D-2D’’, *SI Appendix*, Fig. S4**). Our results suggest that chromatin in the nuclei of the ONL of Chx10-TET animals was less condensed compared to TET mice (**Fig. 2E**). All these abnormalities in the structure of TET-deficient photoreceptors led to the fact that even 1-month-old animals were already functionally blind according to ERG tests (***SI Appendix*, Fig. S5**). We would also like to note that Chx10-cre animals exhibit mosaicism in the expression of cre recombinase in RPCs, which leads to the fact that the effect of TET inactivation can be stronger or weaker in different parts of the retina(18). We found this effect during TEM examination. While in most Chx10-TET retinas the OS and synapses were poorly developed or absent, small areas could be found where the OS and synapses were well developed (***SI Appendix*, Fig. S4**). This could explain the presence of weak photoreceptor activity in Chx10-TET animals assayed by ERG.

### Genetic ablation of the TET family has less impact on the development of non-photoreceptors in the retina

We also assessed in our study the effect of inactivation of the TET-dependent DNA demethylation pathway on other retinal cell types. In addition to significant changes in photoreceptor morphology, we found some changes in Muller glia morphology (**Fig. 3A-3D**). We found that expression of a Muller glia marker Glul was decreased using western blot analysis (**Fig. 3E**). We did not detect significant changes associated with bipolar, amacrine, and horizontal cells (**Fig. 3C, 3F**). However, synaptic contacts in the IPL were less structured in Chx10-TET retinas (**Fig. 3F**). We also measured the thickness of the animals’ optic nerve at 0.5 mm, 2 mm, and 3.5 mm from the globe, and then averaged these values. We found that the optic nerves in Chx10-TET mice were thinner than in TET mice (260±6 µm vs. 357±5 µm, P value < 0.0001, n=10-12 optic nerves; **Fig. 3G**). Since the axons of RGCs form the optic nerve, we analyzed the density of these neurons in the ganglion cell layer (GCL). We found that density was lower in Chx10-TET compared to TET mice (RGC per mm^2^: 3044±109 vs. 3557±91, P value < 0.01, n=5; **Fig. 3H**). The retinal area was also smaller in Chx10-TET mice (13.2±0.9 vs. 16.3±0.2 mm^2^, P value < 0.01, n=5; **Fig. 3I**). A smaller retinal area and RGC density result in a smaller total number of RGCs and, as a result, a smaller number of axons and, accordingly, a smaller thickness of the Chx10-TET optic nerves (**Fig. 3G, 3J**).

**Fig. 3.**
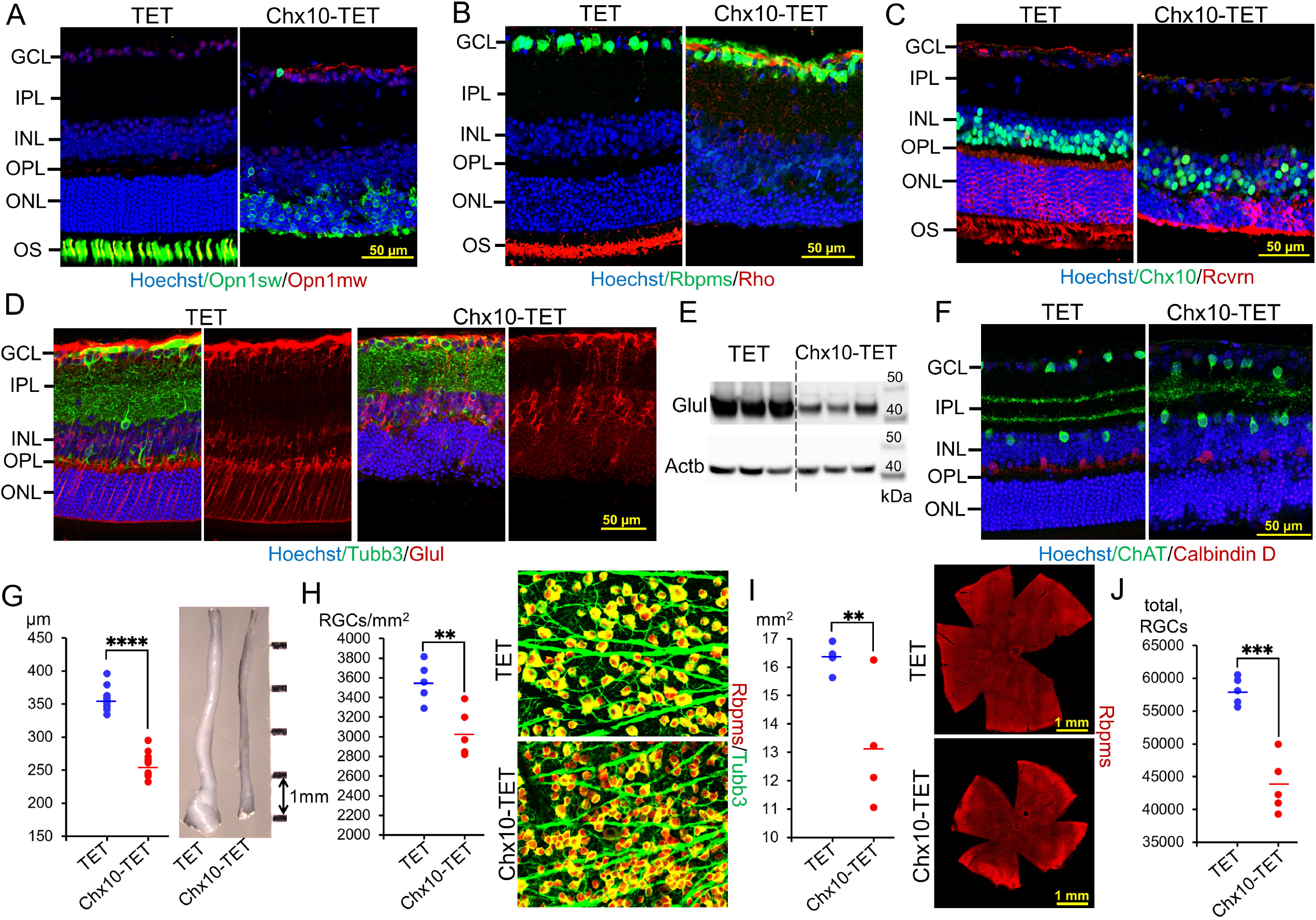
The TET-dependent DNA demethylation pathway has less influence on the differentiation of non-photoreceptors from RPCs. **(A, B)** While s-opsin (Opn1sw is a marker of blue cones) can still be detected in some parts of the retinas in 3-month-old Chx10-TET mice **(A)**, m-opsin (Opn1mw **[A]** is a marker of green cones) and rhodopsin (Rho **[B]** is a marker of rods) are absent in the retinas of Chx10-TET mice. **C)** Rcvrn, as a marker of photoreceptors and bipolar cells, was less expressed in the Chx10-TET retinas. However, bipolar cells (Chx10 as a marker) were present in the retinas of Chx10-TET animals. **(D, E)** Muller glia (Glul as a marker) was less organized in the retinas of 3-month-old Chx10-TET mice **(D)**. The expression of the marker of these cells was reduced as follows from the western blot. **F)** Horizontal cells (Calbindin D as a marker) and amacrine cells (ChAT as a marker) were present in Chx10-TET retinas. However, synaptic contacts were disorganized in the retinal IPL of Chx10-TET animals. **G)** The thickness of the optic nerves was significantly less in 3-month-old Chx10-TET than in TET mice (n=10-12 optic nerves or 5-6 mice per group; P value <0.0001 [^****^]). **H)** The density of RGCs in the ganglion cell layer (GCL) of the flatmounted retina was determined by counting Rbpms-positive neurons in different parts of the retina of known area (n=5 retinas per group, 3-month-old mice; P value <0.01 [^**^]). **I)** The area of the entire retina was significantly smaller in 3-month-old Chx10-TET compared to TET mice (n=5 retinas per group; P value <0.01 [^**^]). **J)** The resulting neuron densities and retinal areas allowed us to determine the total number of RGCs in Chx10-TET and TET mice (n=5 retinas per group; P value <0.01 [^**^]).

### Expression of genes required for photoreceptor development and function is significantly reduced in TET-deficient retinas

Photoreceptor developmental disorders upon inactivation of the TET-dependent DNA demethylation pathway should be reflected in the expression of the corresponding genes. To this end, we investigated gene expression in the retinas of Chx10-TET and TET animals using RNA-seq analysis. Since photoreceptors make up 70-80% of all cells in the retina, studying the gene expression of the whole retina allows us to study the gene expression of the photoreceptors(19). The retinas were collected at P14 and one month (1m) and used for preparation of RNA-seq libraries for next-generation sequencing (NGS; P14, n = 3; 1m, n=4). We chose P14 and 1m animals because the ONL thickness differed only slightly in Chx10-TET and TET mice of these ages. We found that gene expression was significantly different between Chx10-TET and TET retinas (**Fig. 4A-4C, *SI Appendix*, Data S1**). We were somewhat surprised by the lack of difference in the expression of *Tet1, Tet2*, and *Tet3*. However, when we considered the expression of only those TET transcripts in which all exons are retained, we found a significant decrease in the expression of these genes (***SI Appendix*, Fig. S6**). These results indicate the stability of the *Tet1, Tet2*, and *Tet3* transcripts even in the absence of several exons in them. We used k-means clustering to identify coherently expressed groups of genes. We found that the cluster in which Chx10-TET vs. TET gene expression is reduced includes many genes responsible for the development and function of photoreceptors (**Fig. 4D**). Overall, our results indicate that the expression of genes required for phototransduction, as well as OS, inner segment, cilium, and synapse development and function, was significantly reduced in Chx10-TET retinas compared to TET (**Fig. 4E, 4F, *SI Appendix*, Data S1**). These results explain the morphological changes we observed in Chx10-TET photoreceptors (**Figs. 1** and **2**). We found significantly reduced expression of 56 genes, each of which leads to a form of either retinitis pigmentosa, cone or cone-rod dystrophy, congenital stationary night blindness, or Leber congenital amaurosis (**Fig. 4G, *SI Appendix*, Data S1**). Our RNA-seq data agreed with the results of western blot analysis: Nr2e3, Rho, Pde6a, Pde6b, Prph2, Rcvrn proteins were virtually undetectable in Chx10-TET retinas (**Fig. 4H**). Our results also indicate that most of the genes whose expression was decreased are critical for rod photoreceptor development and function (**Fig. 4F, *SI Appendix*, Dataset S1**). Meanwhile, the expression of some genes necessary for cone photoreceptor function was increased (e.g., *Opn1sw, Pde6c, Gnat2, Gngt2, Arr3, Slc24a, Cnga3, Cngb3*, **Fig. 4F, *SI Appendix*, Dataset S1**). These results can be explained by the fact that the expression of *Nrl, Nr2e3, Esrrb*, and *Samd7* transcription factors essential for rod development and function was significantly reduced (**Fig. 4F, *SI Appendix*, Dataset S1**). Thus, the TET-dependent DNA demethylation pathway may play a more important role in the development and function of rods than cones. However, reduced expression of genes (e.g., *Prph2, Rom1, Slc24a1, Rcvrn, Mpp4, Cplx4*) important for the function of both types of photoreceptors leads to the negative impact of inactivation of this pathway on both rods and cones. We also found reduced expression of *Pde6h* and *Opn1mw* cone-related genes.

**Fig. 4.**
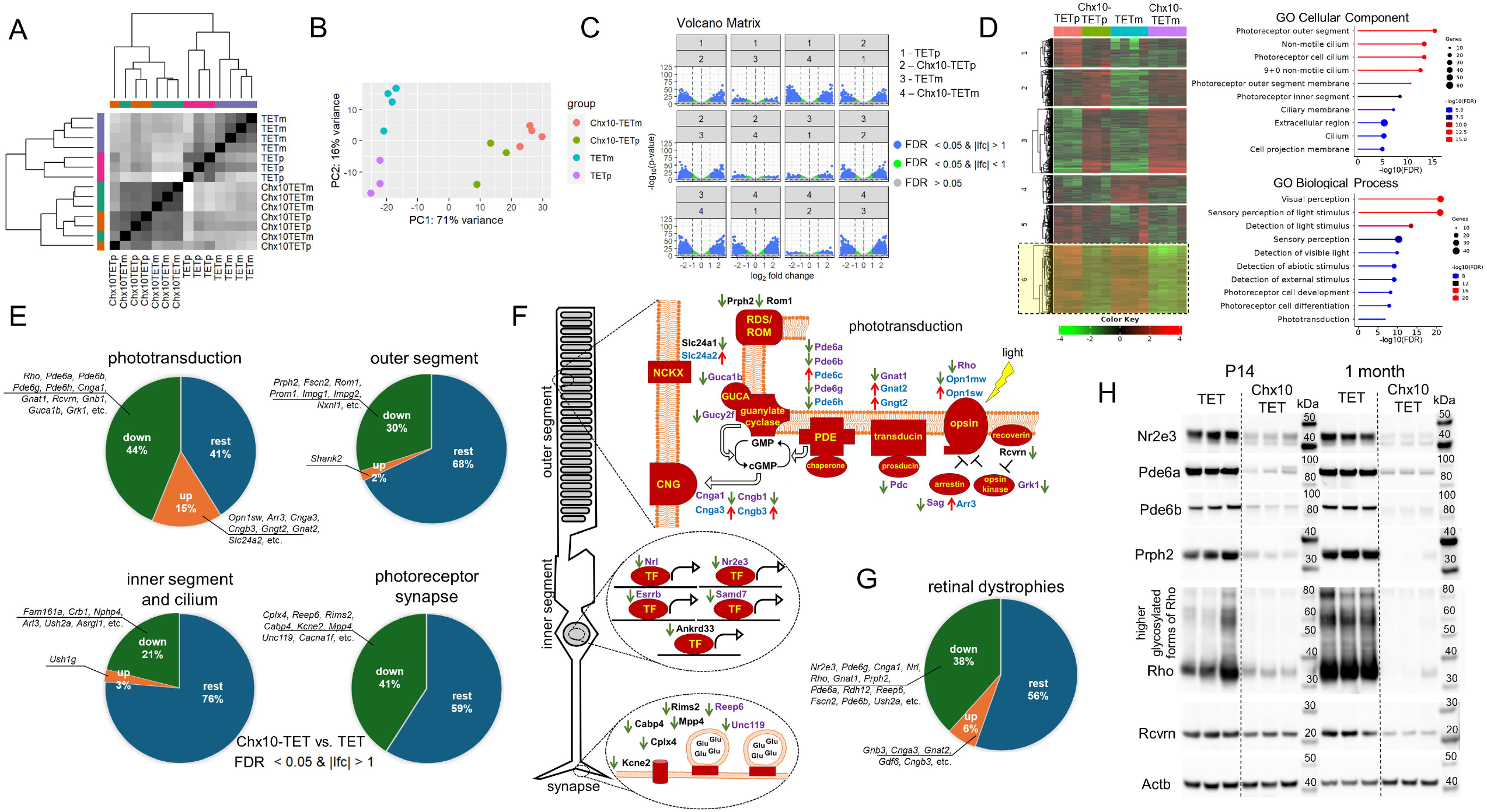
The activity of the TET-dependent DNA demethylation pathway is necessary for the expression of genes responsible for rod specification and for rod and cone maturation. **(A-C)** The sample clustering **(A)**, principal component analysis (PCA, **B**), and volcano plots **(C)** indicate that gene expression in retinas of P14 (p) and 1-month-old (m) Chx10-TET mice differ significantly from gene expression in retinas of TET mice of the same ages. **D)** K-means clustering revealed six distinctive clusters. Cluster six, in which gene expression is significantly reduced in Chx10-TET vs. TET retinas, contains predominantly genes necessary for the development and function of photoreceptors. **(E, F)** The figure **(E)** shows the percentages of genes with decreased (green), increased (orange), and unchanged (dark blue) expression depending on the function they perform in photoreceptors. Many genes whose expression was reduced are responsible for the specification and maturation of rods (**F**, purple). The expression of many genes responsible for cone maturation is increased (**F**, blue). At the same time, the expression of genes necessary for the maturation of both rod and cone photoreceptors is reduced. (**F**, black). **G)** Mutations in many genes whose expression has been reduced (green) in Chx10-TET vs. TET retinas lead to various types of retinal dystrophies. **H)** The reduced expression of *Nr2e3, Pde6a, Pde6b, Prph2, Rho*, and *Rcvrn* genes is consistent with the low levels of the proteins they encode in the retinas of the Chx10-TET animals, as follows from the western blot analysis. Anti-β-actin (anti-Actb) antibody was used to control the loading.

### Genetic ablation of the TET family prevents demethylation of the promoters of genes essential for photoreceptor development and function during the differentiation of RPCs into rods and cones

Disturbances in the activity of TET enzymes should lead to changes in DNA methylation patterns. To study methylation patterns in retinas of Chx10-TET and TET animals, we used whole genome bisulfite sequencing (WGBS). We used whole P14 retinas in our study because 70-80% of their cells are photoreceptors(19). Thus, changes in methylation of photoreceptor gene promoters can be easily detected. The DNA samples isolated from retinas (Chx10-TET, n=3 and TET, n=3) were used in the preparation of six WGBS libraries for NGS. To analyze our NGS data, we used methylKit Bioconductor R package (20). The CpG methylation clustering and principal component analysis indicate a clear difference between the methylation patterns in DNA isolated from the retinas of Chx10-TET vs. TET mice (**Fig. 5A, 5B, *SI Appendix*, Dataset S2**). By combining the capabilities of methylKit and annotatr Bioconductor R packages, we formed a list of genes in which all promoters and first exons were hypermethylated in the retinas of Chx10-TET animals; methylation of at least one promoter or the first exon of these genes should be low in the retinas of TET mice (***SI Appendix*, Dataset S2**)(20, 21). We used this list to analyze the biological processes and diseases in which these genes are involved. We also investigated what cellular components of the proteins encoded by these genes could be a part of. A significant number of genes we found are involved in the development and function of photoreceptors (**Fig. 5C-5E, *SI Appendix*, Dataset S2**). Virtually all diseases are various forms of retinal dystrophies (**Fig. 5D**). Analysis of the DNA methylation and expression levels of these genes showed that high levels of methylation are associated with low levels of expression of these genes in Chx10-TET animals (**Fig. 5E**). Conversely, low methylation levels are associated with high expression levels of these genes in TET mice (**Fig. 5E**). We also found significant changes in mean methylation levels and methylation of individual cytosines in Chx10-TET vs. TET mice at locations close to the transcription start sites (TSS) of studied genes (**Fig. 5F, *SI Appendix*, Fig. S7**). These results suggest that high levels of cytosine methylation in TSS may interfere with the efficient initiation of transcription of genes necessary for photoreceptor development and function, causing retinal abnormalities. Our results also indicate that a significant number of genes whose promoters remain hypermethylated in Chx10-TET mouse retinas are important for rod development and function (e.g., *Nr2e3, Rho, Pde6a, Pde6b, Pde6g, Cnga1, Grk1*). Of particular note, there is a high level of methylation of *Nr2e3* transcription factor, without which the proper development of rods is not possible(3, 4). However, high methylation levels of the promoters of genes that are important for function of both photoreceptor types (e.g., *Prph2, Rcvrn, Cplx4*) should have an impact on the function of cones in Chx10-TET mice.

**Fig. 5.**
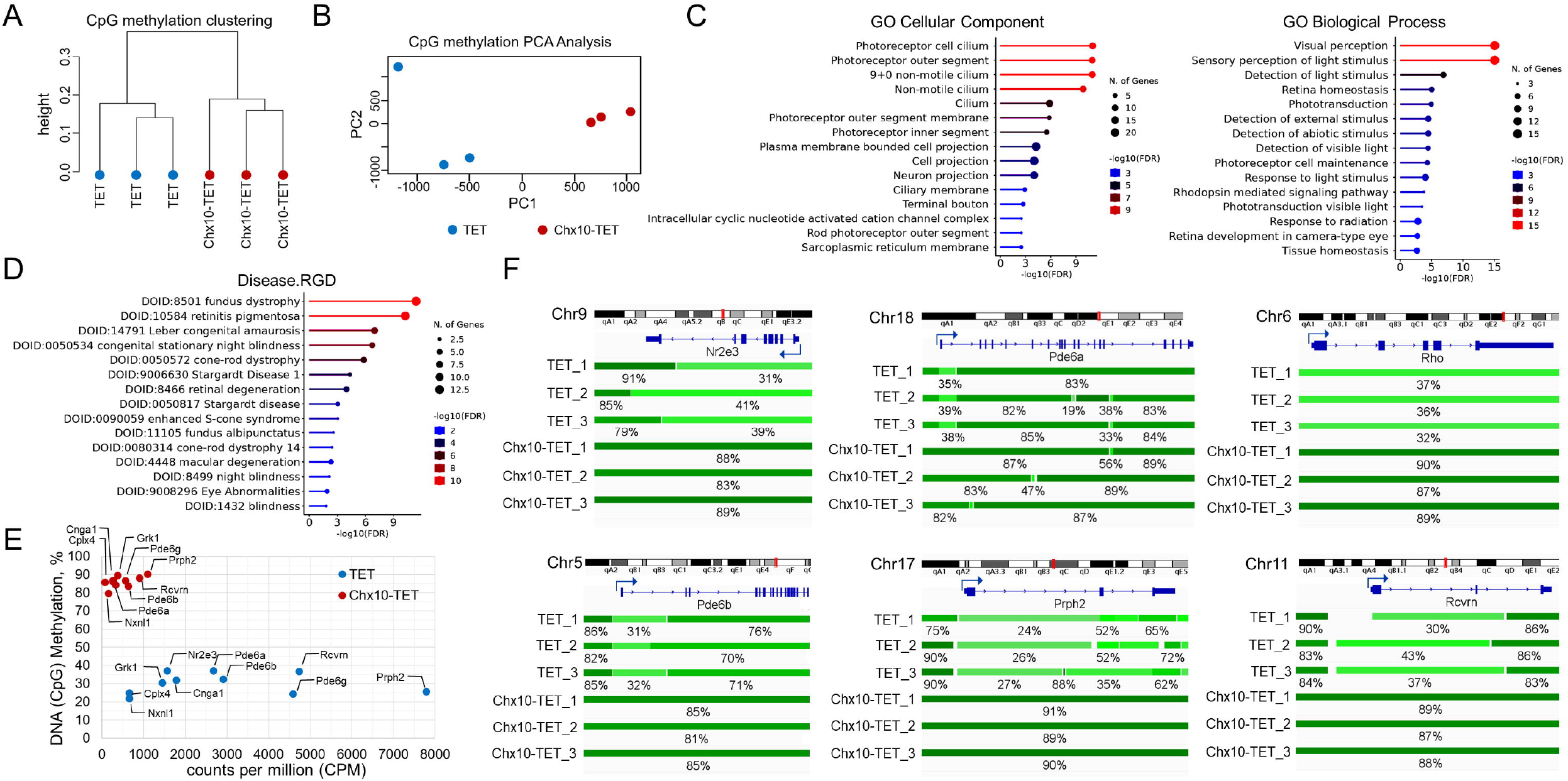
Disruption of the TET-dependent DNA demethylation pathway prevents demethylation of the promoters of genes essential for photoreceptor development and function. **(A, B)** The CpG methylation clustering **(A)** and principal component analysis **(B)** indicate that DNA isolated from retinas of P14 Chx10-TET mice has methylation patterns that are significantly different from that of DNA isolated from retinas of P14 TET mice. (**C, D**) The results of the analysis of genes whose promoters remain hypermethylated in Chx10-TET retinas but are demethylated in TET retinas indicate their predominant association with the development **(C)**, function **(C)**, and pathology **(D)** of photoreceptors. **E)** The figure shows the relationship between the level of promoter methylation and the expression of the corresponding genes. **F)** Visualization of the average % CpG methylation corresponding to different DNA regions indicates that the promoters of genes important for the development and function of photoreceptors were hypermethylated in Chx10-TET retinas and hypomethylated (low methylated) in TET retinas. We used Integrative Genomics Viewer (IGV) to visualize the bed files that were generated by Bismark Bisulfite Mapper.

### TET enzyme inactivation in RPCs significantly reduces DNA hydroxymethylation levels in the retina

The TET enzymes demethylate DNA by oxidizing 5mC into 5-hydroxymethylcytosine (5hmC), 5-formylcytosine (5fC), and 5-carboxylcytosine (5caC)(10, 11). While 5fC and 5caC are quickly replaced by cytosines, 5hmC is stable in nondividing cells and can be detected(10, 11). We assess the effect of TET enzyme inactivation in RPCs on 5hmC levels in the retina. To this end, we used DNA isolated from retinas of P14 Chx10-TET and TET animals and the 5hmC-Seal technology that allows collecting DNA fragments enriched with 5hmC(22). To test specificity and efficiency of the 5hmC enrichment, we used spike-in DNA containing an equimolar mixture of 5mC- and 5hmC-rich DNA fragments. Analysis of our NGS data showed a significant excess of spike-in reads containing 5hmC over 5mC (more than 200-fold for TET and more than 500-fold for Chx10-TET). These results indicated a high level of specificity of our NGS data. We used spike-in normalization to correct for global changes in our NGS data(23). We found significant differences between Chx10-TET and TET samples using clustering and principal component analysis (**Fig. 6A, 6B**). The observed differences between the samples are due to the low levels of 5hmC in the retinas of Chx10-TET compared to TET mice (**Fig. 6C, *SI Appendix*, Dataset S3**). Our data also indicate that 5hmC levels are low in TSS and in CpG islands of TET retinas (**Fig. 6D, 6E**). The patterns of methylation and hydroxymethylation are very similar, which is not surprising since WGBS cannot distinguish 5mC from 5hmC (**Fig. 6E, 6F**). However, the pattern in the TET retinas corresponds to a combination of 5mC and 5hmC, while the pattern in the Chx10-TET retinas is formed only by 5mC, which can affect gene expression. We found that nearly 7,000 genes contain 5hmC-rich regions in their promoters or first exons (***SI Appendix*, Dataset S3**). Most of them could be classified into the group of housekeeping genes based on the GO analysis (**Fig. 6G**). Among these genes, we also found genes essential for photoreceptor development and function (**Fig. 6H**). As we noted in the previous section, mostly unmodified cytosines are present in TSS of photoreceptor genes. Meanwhile, cytosines located at some distance from TSS and considered to be methylated according to WGBS are in fact hydroxymethylated in the retinas of TET animals (**Fig. 6H**). We can assume that TET enzymes complete their work in areas critical for the initiation of gene expression. Cytosines remain hydroxymethylated in areas less important for the onset of gene expression.

**Fig. 6.**
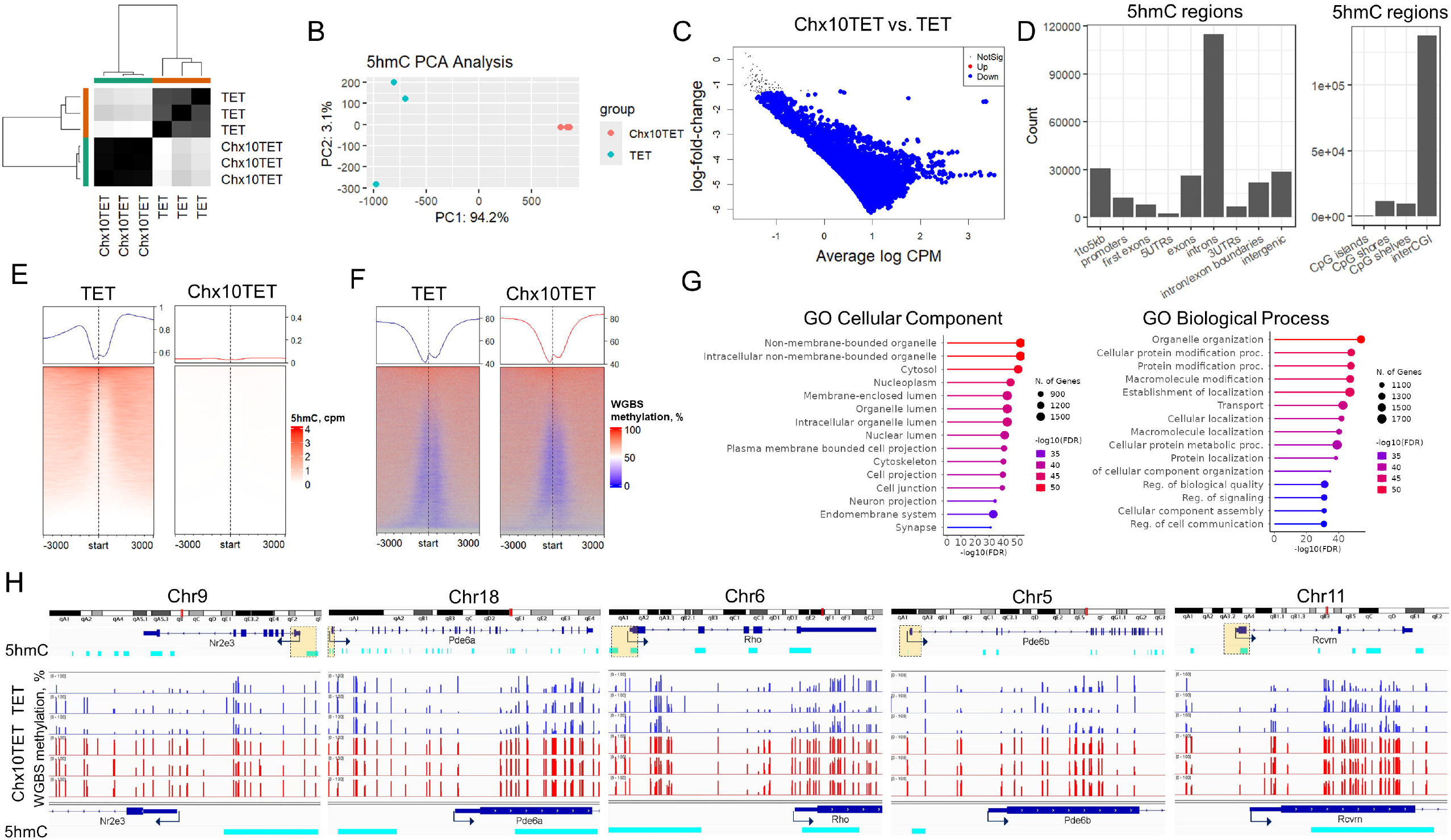
DNA hydroxymethylation levels are reduced in TET-deficient retinas. (**A, B**) DNA hydroxymethylation (5hmC) patterns in TET and Chx10TET retinas differ greatly according to the clustering (**A**) and principal component (PCA, **B**) analysis. **C)** Comparative analysis of 5hmC-rich regions indicates their low content in the retinas of Chx10TET compared to TET animals. The analysis was conducted using the csaw and edgeR R Bioconductor packages (CPM - counts per million). **D)** 5hmC-rich genomic regions are predominantly located in introns, exons, and regions adjacent to promoters (1to5kb). (**E, F**) Hydroxymethylation (**E**) and WGBS methylation (**F**) patterns are similar in the regions containing the transcription start site (TSS) of genes. We use the term WGBS methylation which implies the presence of both types of nucleotides (5mC and 5hmC) in the genome since WGBS cannot distinguish between them. **G)** Genes containing 5hmC-rich genomic regions in the promoter or first exon were subjected to Gene Ontology (GO) enrichment analysis. **H)** Hydroxymethylation and WGBS methylation are absent at the TSS of photoreceptor genes. We used Integrative Genomics Viewer (IGV) to visualize WGBS methylated cytosines (% of C methylation in CpG context) and 5hmC-rich regions (turquoise) adjacent to the TSS.

## Discussion

Disturbances in the development and function of photoreceptors invariably lead to blindness, which can have devastating consequences on a person’s normal life(5-7). Understanding the mechanisms responsible for the development and function of photoreceptors allows us to restore visual function by imparting activity to abnormal photoreceptors or via the regeneration of lost photoreceptors. While significant progress has been made in understanding the signaling cascades that regulate photoreceptor development and function, the contribution of epigenetic mechanisms to these processes is poorly understood (3, 4, 9, 24). The objective of this study was to explore the role of the TET-dependent DNA demethylation pathway in photoreceptor development and function. We found that genetic ablation of the TET family in RPCs prevents demethylation of the promoters of genes essential for rod specification and for rod and cone maturation. Preservation of methylated cytosines in the promoters of these genes significantly reduced their expression in photoreceptors. All these events interfere with the development of photoreceptor OS and synapses, depriving the retina of functioning rod and cone photoreceptors. Over time, the number of photoreceptors decreases in TET-deficient retinas, leading to retinal dystrophy and blindness.

The differentiation of RPCs into photoreceptors includes a specification stage when the photoreceptor precursors decide whether they will be rods or cones, and a maturation stage when the OS, synapses, and other components of the photoreceptors necessary for their function are formed(3, 4). Otx2 and Crx transcription factors are necessary for RPCs to adopt the fate of photoreceptor precursors(3, 4). Onecut1 and Onecut2, in turn, are responsible for cone specification in the presence of Crx(3, 4, 25-27). Meanwhile, Nrl, Nr2e3, and Crx are responsible for rod specification(3, 4, 25-27). Our findings suggest that DNA methylation does not affect the activity of Crx, Onecut1, and Onecut2 and, thus, does not interfere with cone specification (***SI Appendix*, Dataset S1** and **S2**). We also found increased expression of genes essential for cone maturation in Chx10-TET vs. TET retinas (**Fig. 4F**). Meanwhile, the expression of almost all genes necessary for rod specification and maturation was reduced in Chx10-TET vs. TET retinas (**Fig. 4F**). The promoters of genes, such as *Nr2e3, Samd7, Rho, Pde6a, Pde6b, Pde6g, Prph2, Cplx4, Grk1, Cnga1, Rcvrn*, remained hypermethylated and weakly active in the retinas of Chx10-TET animals (***SI Appendix*, Dataset S1** and **S2**). These results indicate that DNA methylation, as an epigenetic mechanism, regulates the specification and maturation of rod photoreceptors, possibly preventing their appearance at an early stage of retinal development. However, the fact that some genes that must be demethylated during RPC differentiation into photoreceptors are used by cones and rods leads to inevitable issues during cone maturation. We also found that the expression of *Opn1mw* and *Pde6h* required for cone maturation was reduced in the retinas of Chx10-TET mice. Thus, the TET-dependent DNA demethylation pathway contributes to cone maturation.

Disruption of the activity of the TET-dependent DNA demethylation pathway leads to reduced expression of many genes, mutations of which lead to one or more forms of either retinitis pigmentosa, cone and cone-rod dystrophy, congenital stationary night blindness, or Leber congenital amaurosis (**Fig. 4G, *SI Appendix*, Dataset S1**). Thus, it is not surprising that we observed such serious disturbances in the development and function of TET-deficient photoreceptors leading to their death. However, mutations in either *TET1, TET2*, or *TET3* leading to retinal dystrophies have not been detected, possibly due to the functional redundancy of these genes(10, 11, 16, 17). Meanwhile, global knockout of all three genes is lethal for embryos(28). Could, then, any retinal dystrophy forms be caused by disturbances in the process of DNA demethylation? TET enzymes need partners (transcription factors with a DNA binding domain) to bind to hypermethylated DNA and to initiate targeted DNA demethylation(29-35). If there are no such transcription factors or their activity is impaired, then the DNA demethylation process cannot be started even in the presence of TET enzymes. Mutations in transcription factors that prevent TET enzymes from binding to promoters can prevent demethylation of one or more genes necessary for the development and function of photoreceptors (e.g., *Nr2e3, Rho, Pde6a, Pde6b, Pde6g, Prph2, Grk1, Cnga1, Cngb1, Nxnl1, Ush2a*), leading to a form of retinal dystrophy. Phenotypically, it may be similar to the form caused by mutations, but this form of retinal dystrophy will be caused by epigenetic mechanisms in this case. Hard work lies ahead to identify partners of TET enzymes that promote demethylation of genes necessary for the development and function of photoreceptors.

Finally, we have shown in our study that disruption of the activity of the TET-dependent DNA demethylation pathway in RPCs impairs the development and function of photoreceptors, leading to retinal dystrophy and blindness. While the TET-dependent DNA demethylation pathway affects the development of both types of photoreceptors, we found a stronger impact of this pathway on the development of rods than cones. Given that rods are necessary for the survival of cones, even a smaller influence of the TET-dependent DNA demethylation pathway on cone development and function will still inevitably lead to cone death(36). However, slow death of undeveloped photoreceptors (significant retinal degeneration was observed in 6-month-old Chx10-TET animals) creates a window of opportunity to remove methylation and trigger the development of these neurons, restoring their function. Thus, the contribution of retina-specific epigenetic mechanisms to the pathogenesis of retinal dystrophies may significantly change current approaches to diagnosing and treating these diseases.

## Materials and Methods

### Ethical approval

All procedures were executed in compliance with the National Institutes of Health (NIH) Guide for the Care and Use of Laboratory Animals and according to the University of Miami Institutional Animal Care and Use Committee (IACUC) approved protocol (Protocol #: 23-058). Animals were euthanized according to the recommendations of the Panel on Euthanasia of the American Veterinary Medical Association (AVMA). All methods were completed and reported in accordance with ARRIVE guidelines.

### Generation of TET conditional knockouts and genotype analysis

To generate TET triple floxed animals, we crossbred Tet1-floxed, Tet2-floxed, and Tet3-floxed animals (these animals are gifts from Dr. Anjana Rao, La Jolla Institute for Immunology). TET mice served as controls. To genotype TET animals, we used primers: a) Tet1 floxed: F: CAG TTC TAT TCA GTA AGT AAG TGT GCC, R: GGT TGT GTT AAA GTG AGT TGC AAG CG (WT/WT – will have 1 band of 198bp, FL/WT – will have 2 bands [198bp and 242bp], FL/FL – will have 1 band of 242bp); b) Tet2 floxed: F: GCC CAA GAA AGC CAA GAC CAA GAA, R: AAG GAG GGG ACT TTT ACC TCT CAG AGC AA (WT/WT – will have 1 band of 650bp, FL/WT – will have 2 bands [650bp and 780bp], FL/FL – will have 1 band of 780bp); c) Tet3 floxed: F: GGA TGT GAG CTA GTT CTC CTA ACT TGA GAG G, R: CCC TGT CTA CTC TAT TCT TGT GTC AGG AGG (WT/WT – will have 1 band of 195 bp, FL/WT – will have 2 bands [195 bp and 349 bp], FL/FL – will have 1 band of 349bp). We selected only floxed alleles. To inactivate TET enzymes in RPCs, we used Chx10-cre mice (strain #:005105, the Jackson Laboratory). To generate triple conditional Chx10-TET knockout animals, we crossed Chx10-cre and TET mice. To genotype Chx10-TET animals, we used TET primers above and primers to detect cre transgene (F: GCG GTC TGG CAG TAA AAA CTA TC, R: GTG AAA CAG CAT TGC TGT CAC TT). Subsequently, we crossed Chx10-TET and TET mice in order to obtain Chx10-TET and TET littermates. We used male and female mice to address sex as a biological variable. Animals were housed under standard conditions. They were given free access to food and water and had a 12-hour light to dark cycle.

### Electroretinogram (ERG) tests

Prior to scotopic ERG recording, mice were dark-adapted for more than 8 hours with free access to food and water. For photopic ERG recordings, mice were adapted to background white light for 10 minutes. For all ERG tests pupils were dilated with 1% tropicamide for 5 minutes. Mice were anesthetized with 80 mg/kg of ketamine and 10 mg/kg of xylazine delivered IP and positioned dorsally in a headholder. The body temperature was kept at 37±0.5°C with a temperature-controlled heating pad. Animals were screened for cataracts or corneal opacities. The Celeris electrodes were lubricated with Systane gel (0.3% Hypromellose) (Alcon, Geneva, Switzerland) prior to placement on the corneas of each animal. The dual function stimulator-electrodes deliver a light stimulus and record neural electrical activity simultaneously. ERG recordings were collected using the Celeris D430 rodent ERG testing system (Diagnosys LLC, MA) paired with the Espion software (V6.64.14; Diagnosys LLC) following the manufacturer’s protocols. Briefly, both eyes of animals were exposed to a series of single white-light flashes. Scotopic tests subjected dark-adapted animals to intensities of 0.01, 0.1, and 1.0 cd.s/m^2^ at a frequency of 1.00 Hz. B-wave amplitudes were measured from the trough of the a-wave to the peak of the subsequent b-wave. The a-wave’s negative trough was analyzed at an intensity of 1.0 cd.s/m^2^. For photopic ERGs, light-adapted animals were given flash stimuli of 3 and 10 cd.s/m^2^. Both scotopic and photopic ERG recordings were sampled at a frequency of 2000 Hz, covering a duration from 50 ms before to 300 ms after the stimulus. Flicker ERG was performed using continuous 6500K white light flashes with an intensity of 3 cd.s/m^2^ at 10 or 30 Hz, against a background of 30 cd/m^2^ 6500K white light. Data for Flicker ERG was acquired at a sample frequency of 2000 Hz, spanning from 10 ms before to 250 ms after the stimulus. Oscillatory potentials were automatically isolated and measured from the ascending limb of the b-wave using Espion software, which also computed all amplitudes. Each trace marker was reviewed, and any errors were manually corrected.

### Necropsy and tissue collection

To collect eyeballs, retinas, and optic nerves, the mice were injected IP with ketamine (80 mg/kg) and xylazine (10 mg/kg). When we confirmed that the animals were under deep anesthesia, we opened the chest cavity and placed a syringe needle into the heart. The syringe needle was attached to a pump to allow phosphate buffered saline (PBS, pH 7.4; #10010023, ThermoFisher Scientific) to circulate through the animal. Eyeballs and optic nerves were then carefully dissected out, taking care not to apply any pressure on them that could influence the results of morphological analysis and other experiments. Optic nerves were dissected from their insertion point behind the globe all the way to the optic chiasm. The size of the eyeballs and optic nerves were determined on Nikon SMZ1270 stereo microscope using Micrometrics SE Premium software. The retinas removed from the eyes were collected in the appropriate buffers for RNA and DNA purification, transmission electron microscopy (TEM), immunohistochemistry, and western blot analysis.

### Study of the thickness of the Chx10-TET and TET retinal layers

The eyeballs were fixed with 4% paraformaldehyde (PFA, in PBS) and were then placed in 30% sucrose (in PBS) for cryoprotection overnight. The next day, the eyeballs were embedded into Cryo-Gel (Leica), frozen in a −80°C freezer, and then, sectioned on a Leica cryostat at 12 µm. We collected only those retinal sections that contained the optic nerve head. The retinal sections were permeabilized with 0.3% Triton X-100 in PBS, washed with PBS, and blocked in a buffer containing 5% donkey serum, 2% BSA and 0.15% Tween-20 in PBS. To stain retinal sections, we added an anti-Rho (1:100, PA5-85608, ThermoFisher Scientific) antibody (to label rod outer segments) and peanut lectin (PNA, L32458, ThermoFisher Scientific) from *Arachis hypogaea* (to label cone outer segments) to the blocking buffer overnight. The next day, the retinal sections were washed with PBS and incubated with secondary fluorescent antibodies (ThermoFisher Scientific). Hoechst 33342 (H3570, ThermoFisher Scientific) was used to stain the cell nuclei. Images were collected and analyzed using Leica STELLARIS confocal microscope and its software. We determined the average thickness of each retinal layer every 200 µm starting from the head of the optic nerve. To this end, we determined the area of the retinal layer between the boundaries (e.g., between 400 µm and 600 µm). We then measured the length between the boundaries in the layer (this length could differ from 200 µm since the retina is curved) and divided the area by this length. We considered the resulting value to be the average thickness.

### Terminal deoxynucleotidyl transferase (TdT) dUTP Nick-End Labeling (TUNEL) assay

To detect dead cells in the retinas, we used the Click-iT Plus TUNEL Assay Kit (C10617, ThermoFisher Scientific) according to manufacturer’s instructions.

### Transmission electron microscopy (TEM)

The collected experimental and control retinas were fixed in 2% glutaraldehyde in 0.05 M phosphate buffer and 100 mM sucrose, post-fixed overnight in 1% osmium tetroxide in 0.1 M phosphate buffer, and then dehydrated through a graded alcohol series. Retinas processed in this way were embedded in a mixture of EM-bed/Araldite (Electron Microscopy Sciences). Sections (1 μm thick) were stained with Richardson’s stain for observation under a light microscope. Ultrathin sections (100 nm thick) were cut with a Leica Ultracut-R ultramicrotome and stained with uranyl acetate and lead citrate. The grids were viewed at 80 kV in a JEOL JEM-1400 (JEOL, Tokyo, Japan) transmission electron microscope and images captured with an AMT BioSprint 12 (AMT Imaging Systems, Woburn, Massachusetts) digital camera.

### Immunohistochemistry

Retinas fixed with 4% PFA were sectioned (100 μm thick) using Leica vibratome. These sections were permeabilized with 0.3% Triton X-100 in PBS, washed with PBS, and blocked in a buffer (5% donkey serum, 2% BSA and 0.15% Tween-20 in PBS). After one-hour, primary antibodies were added to the retinal sections. For these purposes, we used the following antibodies: anti-Rho (1:100, PA5-85608, ThermoFisher Scientific), anti-Opn1sw (1:200, sc-14363, Santa Cruz Biotechnology), anti-Opn1mw (1:200, anti-Opsin red/green, AB5405, MilliporeSigma), anti-Rbpms (1:400, GTX118619, GeneTex), anti-Chx10 (1:200, AB9016, MilliporeSigma), anti-Rcvrn (1:500, AB5585, MilliporeSigma), anti-Tubb3 (1:400; 802001, BioLegend), anti-Glul (1:200, ab73593, Abcam), anti-ChAT (1:200, AB114P, MilliporeSigma), anti-Calbindin D (1:300, C9848, MilliporeSigma). The retinal sections were incubated overnight with primary antibodies, and then, the next day, they were washed with PBS, and incubated with species-specific secondary fluorescent antibodies (ThermoFisher Scientific). Control sections were incubated without primary antibodies. We used Hoechst 33342 (H3570, ThermoFisher Scientific) to stain the cell nuclei. Images were collected using Leica STELLARIS confocal microscope. To perform immunohistochemistry of flat-mounted retinas, retinas fixed with PFA were permeabilized with 0.5% Triton X-100 in PBS, blocked with 0.5% Triton X-100 containing 10% donkey serum in PBS, and then incubated overnight in a buffer (0.2% Triton X-100, 10% donkey serum in PBS) containing primary antibodies. The next day, the retinas were washed with PBS, incubated with species-specific secondary fluorescent antibodies, and washed again. Retinas were flatmounted, coverslipped, and imaged with Leica STELLARIS confocal microscope. To determine the density of RGCs in the GCL, the numbers of Rbpms-positive neurons were counted in 20 random retinal images that corresponded to an area of 343µm × 343µm, using ImageJ software (https://imagej.nih.gov/). The retinal area was determined using Leica STELLARIS confocal microscope software.

### RNA-seq library preparation, sequencing, and data analysis

Total RNA was isolated from retinas using RNeasy Plus Mini Kit (#74134, Qiagen). We evaluated RNA quality and measured RNA quantity using 2100 Bioanalyzer Instrument (Agilent Technologies), Qubit 4 Fluorometer, and the NanoDrop One spectrophotometer (both from ThermoFisher Scientific). We used RNA samples with a RIN score of 8 or higher. To prepare non-stranded RNA-seq libraries, we used mRNA-seq Lib Prep Kit for Illumina (RK20302, ABclonal) according to manufacturer’s instructions. Briefly, poly-T oligo-attached magnetic beads were used to isolate the mRNA from total RNA. After fragmentation, the first strand complementary DNA (cDNA) was produced using random hexamer primers followed by the second strand cDNA synthesis. After end repair, adapter ligation, size selection, amplification, and purification, the RNA-seq libraries were ready for next-generation sequencing (NGS). The RNA-seq libraries were sequenced using 2 × 150 paired end (PE) configuration. The FASTQ files obtained in this study were uploaded to the BioProject database (https://www.ncbi.nlm.nih.gov/bioproject/) and are available under the accession number PRJNA1121391. The STAR RNA-seq aligner, HTseq package, and a basic workflow were applied to calculate how many reads overlap each of the mouse genes (37, 38). To this end, we used GENCODE M25 (GRCm38.p6) Mus musculus reference genome mm10. The differential gene expression analysis was performed using the edgeR R Bioconductor package (39). We used the DESeq2 R Bioconductor package to perform the sample clustering and principal component analysis (PCA)(40). The ViDGER R Bioconductor package and Integrative Genomics Viewer (IGV, https://igv.org/) were applied to visualize the RNA-seq data. We used iDEP (ver. 2.01, http://bioinformatics.sdstate.edu/idep/) for k-means clustering and identification of important signaling cascades and biological processes (41).

### Western blot analysis

Each individual retina was placed in 50 µL of T-PER buffer (#78510, ThermoFisher Scientific) supplemented with protease inhibitor. Total protein concentrations were measured using the Micro BCA Protein Assay Kit (#23235, ThermoFisher Scientific). The protein concentration determined for each individual retina varied from 4µg/µL to 5µg/µL. The SDS-PAGE gel (NuPAGE Bis-Tris Mini Protein Gels, 4–12%, NP0323BOX, ThermoFisher Scientific) wells were loaded with the same amount of protein. The protein samples were resolved on the gel and transferred to PVDF transfer membrane (#88585, ThermoFisher Scientific). The membranes (blots) were blocked in TBST buffer (Tris-buffered saline and 0.15% Tween-20) containing 5% nonfat dry milk (#9999, Cell Signaling Technology). The blots were probed overnight with primary antibodies diluted in blocking buffer. We used the following antibodies: anti-Rho (1:1500, PA5-85608, ThermoFisher Scientific), anti-Pde6b (1:500, PA1-722, ThermoFisher Scientific), anti-Glul (1:1000, ab73593, Abcam), anti-Rcvrn (1:3000, AB5585, MilliporeSigma), anti-Prph2 (1:2000, 18109-1-AP, Proteintech), anti-Pde6a (1:1000, A7915, ABclonal), anti-Nr2e3 (1:2000, 14246-1-AP, Proteintech). The next day, the blots were washed with TBST and then incubated with secondary antibody (1:10 000, Amersham Biosciences) diluted in TBST containing 2.5% nonfat dry milk. Anti-β-actin (anti-Actb) antibody (1:3000; GTX637675, GeneTex) was used to control the loading. Proteins were detected using SuperSignal West Femto Maximum Sensitivity Substrate (#34094, ThermoFisher Scientific) and the ImageQuant LAS 4000 (GE Healthcare).

### Whole genome bisulfite sequencing (WGBS) and data analysis

Genomic DNA was isolated from retinas using the QIAamp DNA Mini Kit (#51304, Qiagen). The DNA concentration was measured using Qubit 4 Fluorometer and the NanoDrop One spectrophotometer. The quality of genomic DNA was assessed using the Agilent fragment analyzer system. Prior to WGBS library preparation, genomic DNA spiked with unmethylated lambda DNA was fragmented via sonication to produce 350 bp DNA fragments using Covaris S220. To prepare WGBS libraries, we used the EZ DNA Methylation-Gold Kit (D5005, Zymo Research) and xGen Methyl-Seq DNA Library Prep Kit (#10009860, Integrated DNA Technologies, Inc.) according to manufacturer’s instructions. WGBS libraries were sequenced from both ends using a 2 × 150 paired end (PE) configuration. The FASTQ files obtained in this study were uploaded to the BioProject database (https://www.ncbi.nlm.nih.gov/bioproject/) and are available under the accession number PRJNA1121391. We used bismark aligner, GENCODE M25 (GRCm38.p6) Mus musculus reference (mm10) genome, and a basic workflow to align paired-end reads (42). To determine the methylation state of cytosines and to compute the percentage of methylation, we used the cytosine2coverage and bismark_methylation_extractor modules of bismark. To get the average X coverage, we used the SAMtools software and our bam files. The results of our calculations indicate that the average base coverage of the genome was 11.4X for TET_1, 9.4X for TET_2, 10.1X for TET_3, 9.2X for Chx10-TET_1, 11.7X for Chx10-TET_2, 15X for Chx10-TET_3. Thus, the total average base coverage of the TET genome was 30.9X, and the total average base coverage of the Chx10-TET genome was 35.9X. CpG methylation clustering, CPG methylation PCA analysis, average DNA methylation levels, and identification of genomic region segmentation classes were accomplished using the methylKit R Bioconductor package (20). Annotation was performed using the annotatr R Bioconductor package (21). We used ShinyGO (ver. 0.80, http://bioinformatics.sdstate.edu/go/) to identify important signaling cascades and biological processes. The Integrative Genomics Viewer (IGV, https://igv.org/) was used to visualize methylated cytosines.

### 5hmC-Seal NGS library preparation and analysis

Genomic DNA was isolated, and its quality and quantity were assessed as described above. DNA was fragmented via sonication to produce 200 bp DNA fragments using Covaris M220. Prior to the 5hmC enrichment, fragmented DNA was spike-in with an equimolar mixture of methylated (5mC) and hydroxymethylated (5hmC) different double strand DNA (dsDNA) fragments from the rabbit IgH locus (#55023, Active Motif). To capture DNA fragments containing 5hmC, we used the 5hmC Profiling Kit (#55023, Active Motif) based on the 5hmC-Seal technology. To prepare NGS libraries, we used NEBNext Ultra II DNA Library Prep Kit for Illumina (E7645S, New England Biolabs) and NEBNext Multiplex Oligos for Illumina (96 Unique Dual Index Primer Pairs, E6440S, New England Biolabs) according to manufacturer’s instructions. 5hmC-Seal NGS libraries were sequenced from both ends using a 2 × 150 PE configuration using the NovaSeq X Plus system (Illumina). The FASTQ files were uploaded to the BioProject database (https://www.ncbi.nlm.nih.gov/bioproject/) and are available under the accession number PRJNA1121391. We used FastQC to evaluate the quality of each FASTQ file. We used the Bowtie2 tool to align paired-end reads using default parameters(43). To create the Bowtie2 index for the reference genome, we used GENCODE M25 (GRCm38.p6) Mus musculus reference (mm10) genome and the two spike-in DNA fragment sequences (refer to the Active Motif 5hmC Profiling Kit manual). We found that more than 80% of reads were aligned concordantly exactly 1 time. We used the SAMtools software to obtain sorted BAM files from SAM files. 5hmC-enriched genomic regions were called with csaw R Bioconductor package with flags: minq=10, max.frag=600, discard=blacklist, pe=“both”, dedup=FALSE, width=50, filter=30(44). Reads overlapping blacklisted regions (ENCODE Blacklist Regions) were removed with discard=blacklist flag. Low-abundance windows were filtered and removed by computing the coverage in each window compared to a global estimate of background enrichment with flags: bin=TRUE, width=10000. We applied a fold-change threshold > 5. In order to spike-in normalize the data and subsequently identify enriched regions in TET retinas compared to Chx10-TET retinas using the edgeR R Bioconductor package, deduplication was not performed (dedup=FALSE). Spike-in normalization was accomplished by equalizing the number of reads corresponding to 5hmC dsDNA 207 bp fragment from the rabbit IgH locus between all libraries (#55023, Active Motif). Normalization factors were computed using the TMM method and normFactors() csaw function. Windows less than 150 bp apart were clustered into 5hmC regions using mergeResults() csaw function. We used dist, heatmap.2, prcomp and other R functions for clustering and PCA analysis. We used the annotatr R Bioconductor package to characterize 5hmC-enriched genomic regions. To visualize the enrichment of 5hmC and 5mC in genomic regions we used the EnrichedHeatmap R Bioconductor package with flags: extend = 3000, w = 50, mean_mode = “w0” (for 5hmC data), mean_mode = “absolute” (for WGBS data)(45). To this end, we used unfiltered normalized csaw data. We used ShinyGO for gene ontology enrichment (GO) analysis. The Integrative Genomics Viewer (IGV) was used to visualize 5hmC regions and 5mC.

## Statistical Analysis

The unpaired Student’s t-test was applied for experiments containing one variable. For experiments containing two or more variables, a one-way analysis of variance (ANOVA) was utilized. P values equal to or less than 0.05 were considered statistically significant. Negative and positive controls were always used in our study. Animal experiments utilized offspring from several different breeding pairs in every experimental group to avoid the potential for a unique genetic bias. Generation and analysis of next-generation sequencing (NGS) data were carried out in-house according to ENCODE standards and pipelines. All statistics were performed on Prism GraphPad 10.1.1, with R (version 4.4.0), or Microsoft Excel.

## Data availability

The datasets collected and analyzed in this study are available in the BioProject database (accession number PRJNA1121391) and in the article/Supplementary Data. We also used Gene Expression Omnibus (GEO) NCBI GSE101986 and GSE104827 datasets to examine the expression of *Tet1, Tet2*, and *Tet3* genes in developing mouse and human retinas.

## Code availability

This study did not use any unique custom code, algorithms or software. We used only the open-source software specified in the article.

## Acknowledgments

This study was supported in part by the National Institutes of Health/National Eye Institute, grants R01 EY035235 (D.I.), National Institutes of Health/National Cancer Institute R01 CA248890 (D.P.), National Institutes of Health/National Eye Institute Center Core grant P30 EY014801, Research to Prevent Blindness/Unrestricted Grant GR004596-1, and the Mark J. Daily Inherited Retinal Diseases Research Center at Bascom Palmer Eye Institute. We are grateful to Dr. Anjana Rao (La Jolla Institute for Immunology) for the animals she provided to us. The authors thank Charles K. Yaros for his expert assistance. We acknowledge the University of Miami Transmission Electron Microscopy Core for EM sample preparation and assistance with the generation of EM images.

