## Supplementary material for "Genetic ablation of the TET family in retinal progenitor cells impairs photoreceptor development and leads to blindness": SI Appendix

**This PDF file includes:**

Supporting text
Figures S1 to S7
Legends for Datasets S1 to S3

**Other supporting materials for this manuscript include the following:**

Datasets S1 to S3

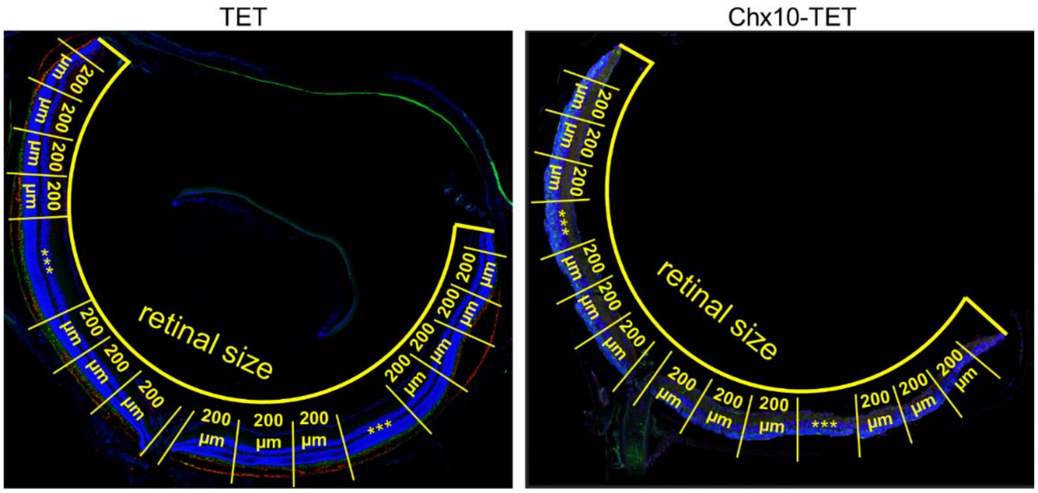


**Fig. S1.** The thickness of the TET and Chx10-TET retinal layers was measured every 200 µm, starting from the head of the optic nerve. The retinal size meant the length of the arc passing from the periphery of the retina on the left through the optic nerve head up to the periphery of the retina on the right.

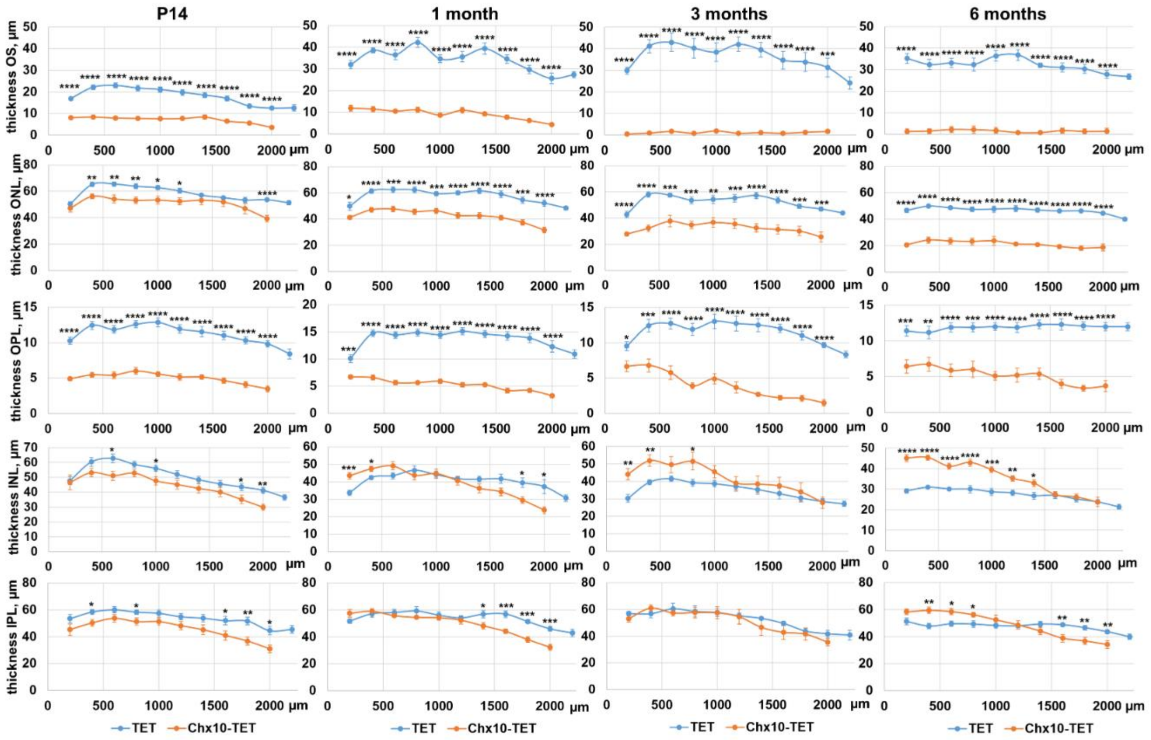


**Fig. S2.** We found that already in postnatal day 14 (P14) Chx10-TET mice, the thickness of the retinal outer segments (OS) and outer plexiform layers (OPL) measured every 200 µm was significantly less compared to the thickness of the OS and OPL of the TET retinas. The difference in the thickness of the outer nuclear layer (ONL) of the Chx10-TET vs. TET retinas was evident in 1-month-old animals and became significant in 3- and 6-month-old animals. In general, the inner nuclear layers (INL) and inner plexiform layers (IPL) did not differ much in animals of all ages studied. The observed increase in the thickness of the INL of the 6-month-old Chx10-TET retinas is likely due to the fact that their cells move and begin to fill the space previously occupied by the cells of the ONL.

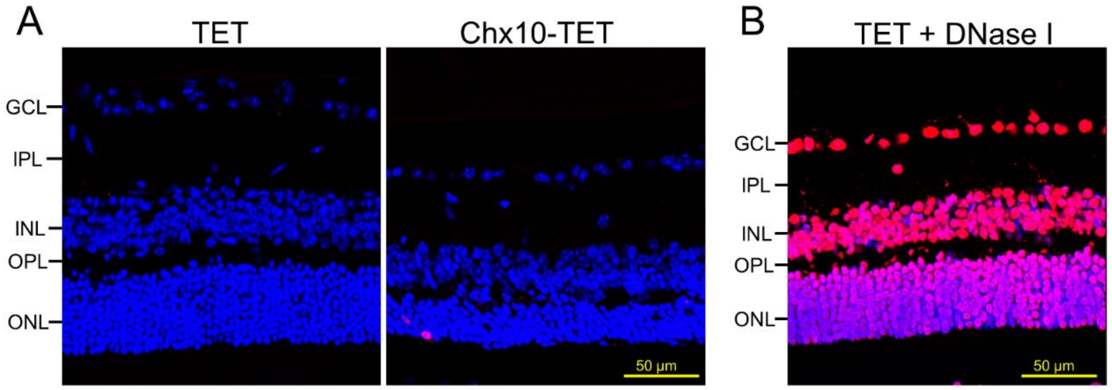


**Fig. S3.** Slow Chx10-TET retinal degeneration occurs because a small number of TET-deficient photoreceptors die over a period of time. **A)** While we did not find TUNEL-positive dead cells in TET retinas, we found only a few TUNEL-positive dead cells in the ONL of Chx10-TET retinas. **B)** Positive controls were treated with DNase I to induce TUNEL-positive DNA strand breaks. (red - TUNEL-positive dead cells, blue - Hoechst 33342-positive cell nuclei)

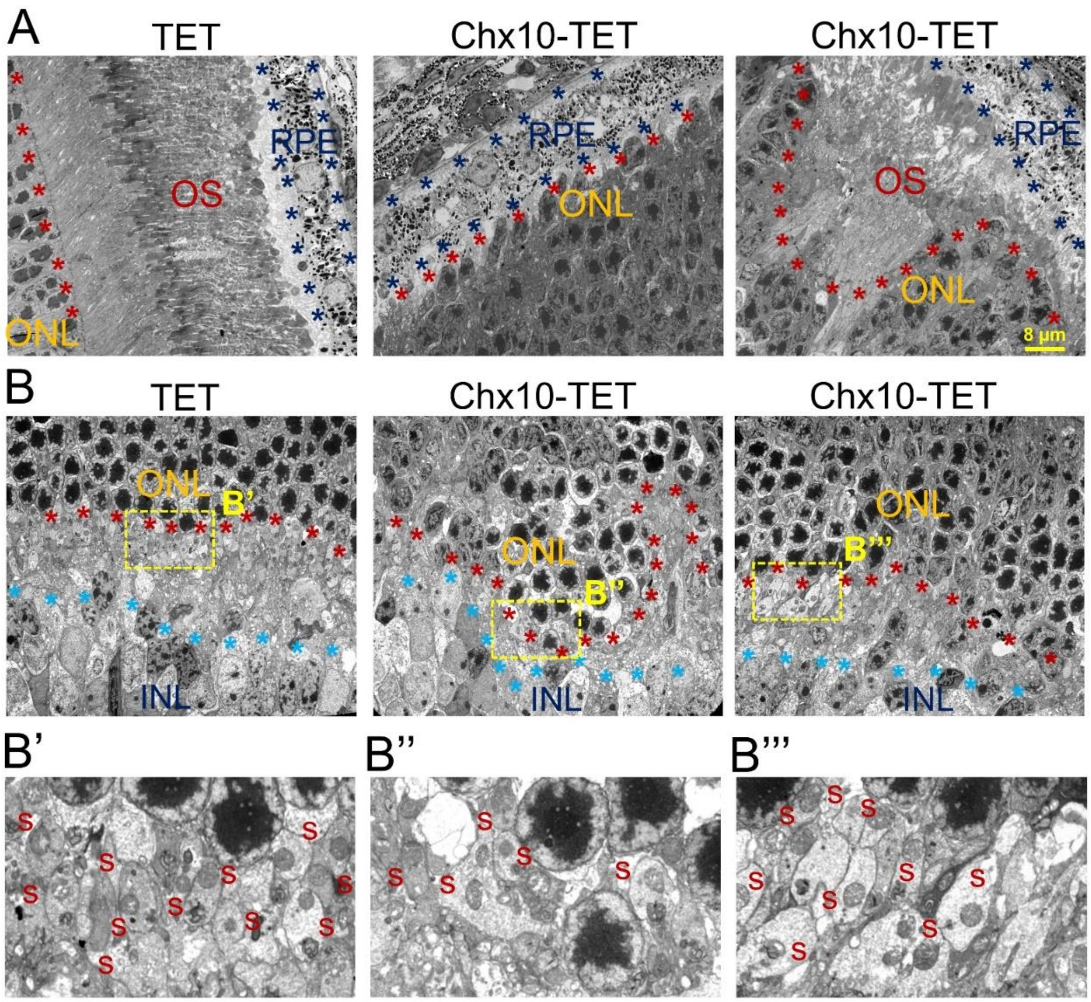


**Fig. S4.** The TET-dependent DNA demethylation pathway is required for the development of photoreceptor outer segments and synapses. **A)** While the OS of the retinas was fully developed in 1-month-old TET animals, transmission electron microscopy (TEM) examination revealed the absence or underdevelopment of the OS in the retinas of 1-month-old Chx10-TET mice. However, due to the mosaic nature of cre recombinase expression in Chx10-cre mice, it was sometimes possible to find relatively well-developed OS in certain areas of the retinas of Chx10-TET animals. **B)** The retinal ONL and INL of 1-month-old TET animals were separated by OPL in which many well-developed synapses (s, B') were identified. The OPL and the synapses in it were difficult to detect in the retinas of 1-month-old Chx10-TET animals (B''). The lack of a barrier between the ONL and INL leads to mixing of their cells in Chx10-TET retinas. However, due to the mosaic nature of cre recombinase expression, it was possible to find areas where synapses were well developed in Chx10-TET animals (B'''). We collected retinas from five 1-month-old TET and Chx10-TET mice for TEM analysis. All images were collected at the same distance from the optic nerve head (800 µm). The dark blue stars limit the layer that contains retinal pigment epithelial (RPE) cells. Red stars indicate the boundary of the outer nuclear layer (ONL). Blue stars indicate the boundary of the inner nuclear layer (INL).

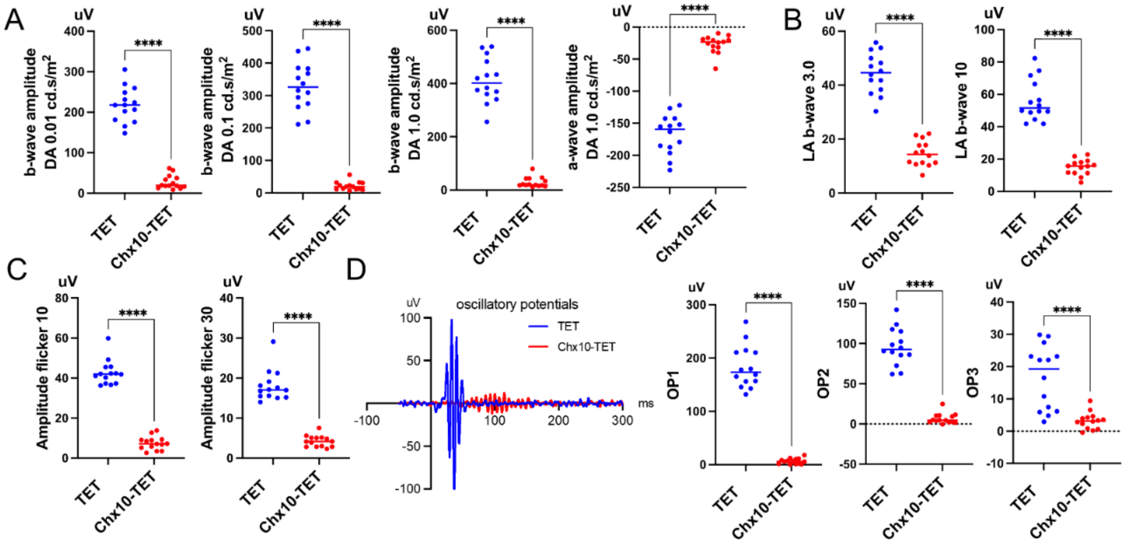

**Fig. S5.** One-month-old Chx10-TET mice were functionally blind as indicated by ERG tests. **A)** Scotopic ERGs: both eyes of dark-adapted animals were subjected to an intensity series of single white-light flash stimulus at 0.01, 0.1, and 1.0 cd.s/m$^2$ at a frequency of 1.00 Hz. B-wave amplitudes (derived primarily from Muller cells and ON-bipolar cells) were analyzed as the amplitude from the trough of the a-wave to the peak to the subsequent b-wave peak. The negative trough of the a-wave (derived from photoreceptors, rods and cones) was analyzed at 1.0 cd.s/m$^2$ intensity. **B)** Photopic ERGs: light-adapted animals were presented with a 3 and 10 cd.s/m2 flash intensity ERG. **C)** Flicker ERG: measures cone opsin regeneration rate. Animals were exposed to continuous 6500K white light flash cycles at an intensity of 3 cd.s/m$^2$ with frequencies of 10 or 30 Hz, against a background of 30 cd/m$^2$ 6500K white light. Flicker amplitudes were measured from the N1 trough to the subsequent P1 peak. **D)** Oscillatory potentials (primarily from amacrine cells) were automatically isolated and measured from the ascending limb of the b-wave by the Epsion software (V6.64.14; Diagnosys LLC) at 1 cd.s/m$^2$. Representative traces are shown. For all tests n=14 eyes (7 animals per group, TET or Chx10-TET) were sampled. For all tests an unpaired t-test was performed between the two groups and was significant to P value <0.0001 (****).

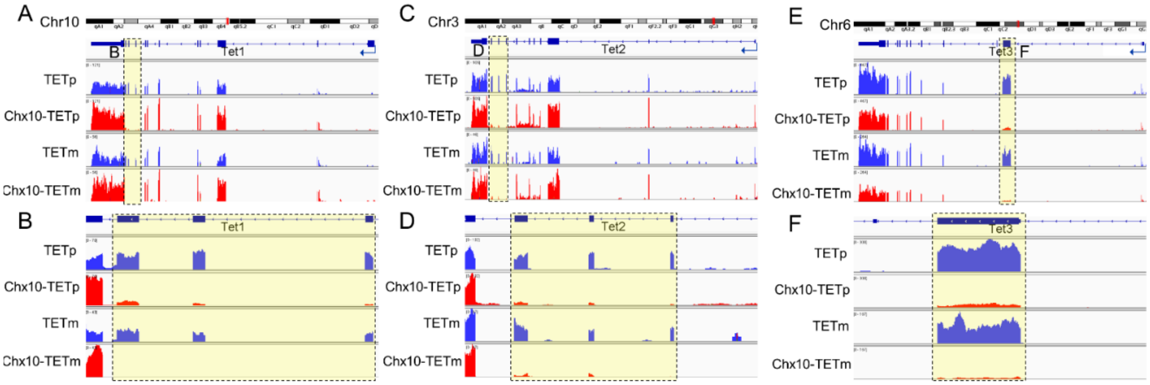


G

| gene | logFC2vs1 | logFC3vs1 | logFC4vs1 | logCPM | F | PValue | FDR |
|---|---|---|---|---|---|---|---|
| *Tet1* | 0.63226916 | -0.86402023 | 0.45203328 | 5.48192 | 31.824 | 8.6E-07 | 1.6E-05 |
| Tet1ex10_12 | -1.82971896 | -1.24111157 | -5.39993763 | -0.5994 | 21.165 | 1.1E-05 | 0.00012 |
| *Tet2* | -0.10867619 | -0.8949554 | -0.54421910 | 5.31088 | 7.5611 | 0.00254 | 0.00822 |
| Tet2ex8_10 | -2.22231757 | -0.75915128 | -4.08112108 | 0.53045 | 37.837 | 2.7E-07 | 6.4E-06 |
| Tet3 | -0.6253074 | -0.58931173 | -1.4060100 | 7.59584 | 22.005 | 8.8E-06 | 0.0001 |
| *Tet3ex3* | -2.98061362 | -0.68361617 | -4.82225517 | 4.51728 | 94.069 | 4.9E-10 | 5.6E-08 |

H

| gene | logFC2vs1 | logCPM | F | PValue | FDR |
|---|---|---|---|---|---|
| *Tet1* | 0.6322052 | 5.48187 | 12.947 | 0.00259 | 0.01842 |
| Tet1ex10_12 | -1.8297189 | -0.5994 | 14.9167 | 0.0015 | 0.0123 |
| *Tet2* | -0.1087368 | 5.31084 | 0.23535 | 0.6345 | 0.7858 |
| Tet2ex8_10 | -2.2223175 | 0.53045 | 37.5163 | 1.8E-05 | 0.00043 |
| *Tet3* | -0.6253667 | 7.59579 | 11.3031 | 0.00421 | 0.02633 |
| Tet3ex3 | -2.9806136 | 4.51728 | 89.4105 | 9.3E-08 | 7.6E-06 |

I

| gene | logFC4vs3 | logCPM | F | PValue | FDR |
|---|---|---|---|---|---|
| *Tet1* | 1.3161268 | 5.48187 | 70.09 | 4.5E-07 | 1.2E-05 |
| Tet1ex10_12 | -4.158826 | -0.5994 | 29.3885 | 6.8E-05 | 0.00069 |
| *Tet2* | 0.3507946 | 5.31084 | 3.10992 | 0.0979 | 0.20575 |
| Tet2ex8_10 | -3.321969 | 0.53045 | 61.5634 | 1E-06 | 2.3E-05 |
| *Tet3* | -0.816618 | 7.59579 | 25.3216 | 0.00014 | 0.00124 |
| Tet3ex3 | -4.138639 | 4.51728 | 176.513 | 9E-10 | 1.1E-07 |

1-TETp
2-Chx10-TETp
3-TETm
4-Chx10-TETm

**Fig. S6.** Genetic ablation of the TET family in RPCs results in a significant reduction in the expression of transcripts encoding Tet1, Tet2, and Tet3 catalytic domains in the adult retina. **(A-F)** Visualization of the RNA-seq data using Integrative Genomics Viewer (IGV) revealed low read content in exons encoding catalytic domains of *Tet1* (**A, B**), *Tet2* (**C, D**), and *Tet3* (**E, F**) in Chx10-TET vs. TET mice. **G)** The expression of *Tet1, Tet2*, and *Tet3*, considering transcripts with and without catalytic domains, does not differ in Chx10-TET and TET retinas. **(H, I)** If we only take into account reads corresponding to exons encoding Tet1 (Tet1ex10_12), Tet2 (Tet2ex8_10), and Tet3 (Tet2ex3) catalytic domains, then the expression of transcripts encoding catalytic domains is significantly reduced in the retinas of P14 (**H**) and 1-month-old (**I**) Chx10-TET mice. These results indicate the stability of *Tet1, Tet2*, and *Tet3* transcripts that do not contain exons encoding catalytic domains.

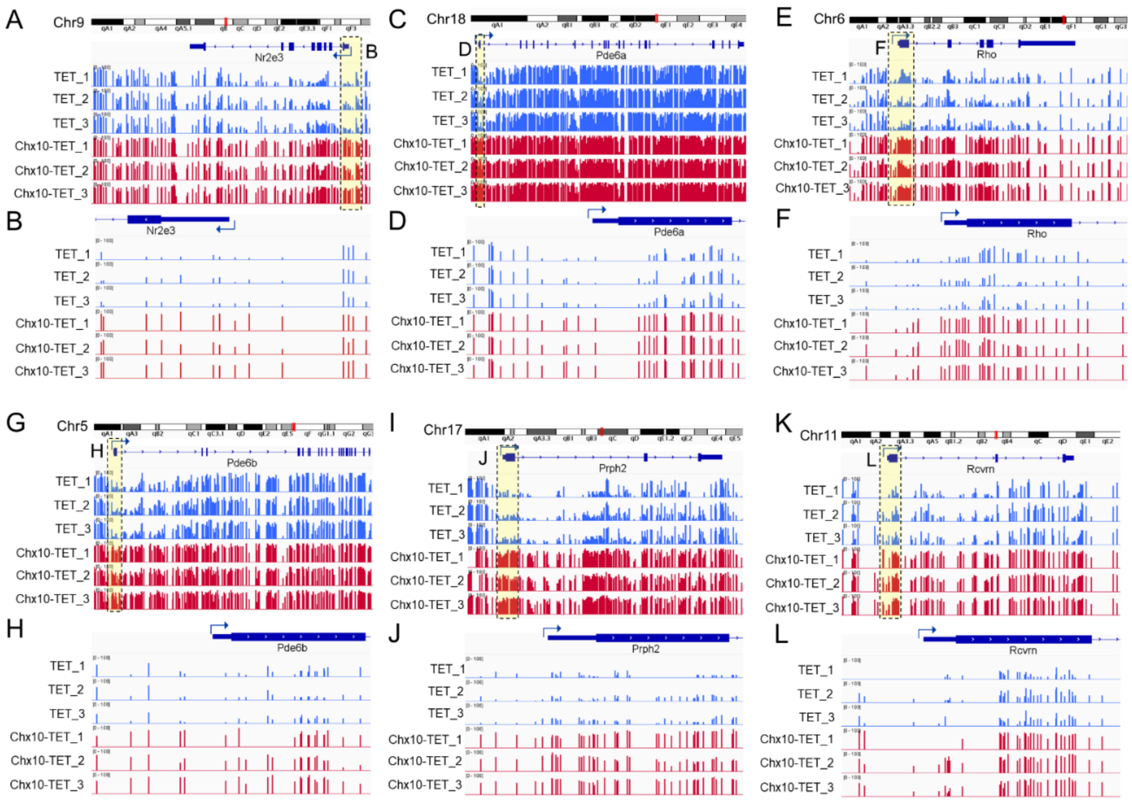


**Fig. S7.** Visualization of individual methylated and unmethylated cytosines in gene promoters indicate a high content of methylated cytosines near the transcription start site (TSS) in DNA isolated from the retinas of P14 Chx10-TET mice. At the same time, cytosines located at the same positions were low methylated (hypomethylated) in DNA isolated from P14 TET mouse retinas. The figure shows the promoters of the following genes: *Nr2e3* (**A, B**), *Pde6a* (**C, D**), *Rho* (**R, F**), *Pde6b* (**G, H**), *Prph2* (**I, J**), and *Rcvrn* (**K, L**). We used Integrative Genomics Viewer (IGV) to visualize the bed files (% of C methylation in CpG context) that were generated by Bismark Bisulfite Mapper.

**Dataset S1 (separate file).** Comparative analysis of gene expression in Chx10-TET and TET retinas: The results of the RNA-seq analysis indicate that the expression of many genes necessary for the development of the photoreceptor outer segment, inner segment, cell cilium, and synapses, as well as the expression of many genes necessary for phototransduction, was significantly reduced in the retinas of P14 and 1-month-old Chx10-TET compared to TET mice. Many of these genes have been implicated in various forms of retinitis pigmentosa, cone and cone-rod dystrophy, congenital stationary night blindness, and Leber congenital amaurosis.

**Dataset S2 (separate file).** Results of analysis of gene methylation levels in Chx10-TET and TET retinas: The combination of methylKit and annotatr Bioconductor R packages made it possible to determine the average level of methylation of promoters and first exons of genes. A search for genes whose promoters and first exons were highly methylated (hypermethylated) in DNA isolated from P14 Chx10-TET retinas compared to P14 TET retinas resulted in a list of genes, many of which are essential for photoreceptor development and function.

**Dataset S3 (separate file).** List of genomic regions enriched with the 5hmC nucleotides: Comparative analysis of genomic regions indicates a low presence of 5hmC in DNA isolated from the retinas of Chx10-TET compared to TET animals. A list of genes that have an increased level of hydroxymethylation in the promoter or first exon is also provided.
